# Comparative morphological and transcriptomic analyses reveal novel chemosensory genes in the poultry red mite, *Dermanyssus gallinae* and knockdown by RNA interference

**DOI:** 10.1101/2020.04.09.034587

**Authors:** Biswajit Bhowmick, Yu Tang, Fang Lin, Øivind Øines, Jianguo Zhao, Chenghong Liao, Rickard Ignell, Bill S. Hansson, Qian Han

**Affiliations:** Laboratory of Tropical Veterinary Medicine and Vector Biology, School of Life and Pharmaceutical Sciences, Hainan University, Haikou, Hainan 570228, China; Norwegian Veterinary Institute, Ullevaalsveien 68 P.boks 750 Sentrum, 0106 Oslo, Norway; Disease Vector Group, Unit of Chemical Ecology, Department of Plant Protection Biology, Swedish University of Agricultural Sciences, Alnarp, Sweden; Department of Evolutionary Neuroethology, Max Planck Institute for Chemical Ecology, 07745, Jena, Germany

**Keywords:** Poultry red mite, *Dermanyssus gallinae*, Sensory morphology, Transcriptome, Chemosensory receptors, Differential gene expression, RNA interference

## Abstract

Detection of chemical cues via chemosensory receptor proteins are essential for most animals, and underlies critical behaviors, including location and discrimination of food resources, identification of sexual partners and avoidance of predators. The current knowledge of how chemical cues are detected is based primarily on data acquired from studies on insects, while our understanding of the molecular basis for chemoreception in acari, mites in particular, remains limited. The poultry red mite (PRM), *Dermanyssus gallinae*, is one of the most important blood-feeding ectoparasites of poultry. Unlike other ectoparasites on animals, PRM feeds mainly at night. During daytime, these animals hide themselves in crevices around the poultry house. The diversity in habitat usage, as well as the demonstrated host finding and avoidance behaviors suggest that PRM relies on their sense of smell to orchestrate complex behavioral decisions. Comparative transcriptome analyses revealed the presence of candidate variant ionotropic receptors (IRs), odorant binding proteins (OBPs), niemann-pick proteins type C2 (NPC2) and sensory neuron membrane proteins (SNMPs). Some of these proteins were highly and differentially expressed in the forelegs of PRM. Rhodopsin-like G protein-coupled receptors (GPCRs) were also identified, while insect-specific odorant receptors (ORs) and odorant co-receptors (ORcos) were not detected. Furthermore, using scanning electron microscopy (SEM), the tarsomeres of all legs pairs were shown to be equipped with sensilla chaetica with or without tip pores, while wall-pored olfactory sensilla chaetica were restricted to the distal-most tarsomeres of the forelegs. Further, using the conserved odorant binding protein (OBP) as a test case, the results showed that RNA interference (RNAi) can be induced in *D. gallinae* chemosensory tissues. This study is the first to describe the presence of chemosensory genes in any *Dermanyssidae* family. It is also the first report of chemosensory gene knockdown by RNAi in any mite species and demonstrate that their diminutive size, less than 1 mm, is not a major impediment when applying gene knockdown approaches. Our findings make a significant step forward in understanding the chemosensory abilities of *D. gallinae*.

## 1. Introduction

Mites are highly diverse arthropods of the subphylum chelicerata, one of the most diverse group in animal radiations (Sharma, 2018). These eight legged arthropods exhibit a tremendous variation in lifestyle, ranging from saprophagous to herbivorous and from predatory to parasitic feeding behaviors (Van Leeuwen and Dermauw, 2016). Some of these species pose a threat to both animal and human health, such as ticks, mites, spiders and scorpions, and may also vector pathogens of importance to animals, insects, plants and humans (Javed et al., 2013). The poultry red mite (PRM), *Dermanyssus gallinae* (de Geer, 1778), is an obligatory hematophagous ectoparasitic mite that infests a wide range of hosts including wild birds, rodents and mammals (George et al., 2015). The PRM is considered the major pest for the poultry industry, with the annual cost in damage and control estimated at €231 million in European countries alone (Flochlay et al., 2017; Bhowmick et al., 2019). Unlike other ectoparasites in animals, PRM sucks blood mainly at night, while they hide themselves in cracks and crevices during daytime so that chickens cannot peck and eat them. Their complex behavior makes them difficult to control with conventional acaricides and other treatment practices (Bhowmick et al., 2019). Hence, an increased understanding of the peripheral olfactory system, including the identification of chemosensory receptors and the ultrastructural characteristics of chemosensory sensilla in PRM, is of great veterinary and economical interest.

Ticks and mites are two different species, but they are evolutionary related groups of arthropods belonging to the class arachnida and the subclass acari. Unlike insects, olfactory organs in mites and other arachnids are located exclusively on the legs (Carr and Roe, 2016). The four pairs of walking legs are covered with specialized hairs, called sensilla, which protect sensory neurons from the insults of the external environment. Three major types of chemosensory sensilla have so far been characterized in arachnids, including the foreleg tarsi of the mites *Varroa destructor* (Dillier et al., 2006; Häußermann et al., 2015), *D. prognephilus* (Davis and Camin, 1976) and *D. gallinae* (Cruz et al., 2005). These unique structures may advantageously allow for the coordination of mechano-, hygro-, thermo and chemoreception (Barth, 2002; Willemart et al., 2007). Even though the identification of structure and the internal arrangement of sensilla is an effective approach to judge their functions (Altner and Prillinger, 1980), very few studies have been conducted in the acarines as a whole, and especially in mites (Tichy and Barth, 1992). Thus, detailed ultrastructural studies using scanning electron microscopy (SEM) would improve our knowledge of poultry red mite sensory biology dramatically.

The vast majority of receptors that detect chemosensory stimuli and convert ligand binding into neural activity belong to one of three membrane-bound receptor families: gustatory receptors (GRs), odorant receptors (ORs) and ionotropic receptors (IRs). While insects make use of all of these receptor families, non-insect arthropods rely on GRs and IRs for their chemical communication (Vosshall and Stocker 2007; Benton, R, 2009; Groh-Lunow et al., 2015). The GRs detect non-volatile compounds, including bitter and sweet compounds, salts and some gustatory pheromones (Vosshall and Stocker 2007; Thorne et al., 2004), but also carbon dioxide in various insect species (Croset et al., 2010). The IRs are involved in olfaction and gustation, but also have non-chemosensory functions, including detection of humidity and temperature (Van and Garrity, 2017). Proteins from other gene families have also been shown to contribute to taste and olfaction. Members of the odorant binding protein (OBP) and chemosensory protein (CSPs) families in insects are highly concentrated in the sensillum lymph, and have been shown to bind odorant molecules (Vogt et al., 1991; Sánchez-Gracia et al., 2009). OBP-like proteins have been well characterized in chelicerata, performing roles analogous to those of insect OBPs and CSPs (Renthal et al., 2016; Iovinella et al., 2016). Besides OBPs and CSPs, another potential chemosensory protein family, Niemann Pick type C2 (NPC2), has been identified in both insects and non-insect arthropods (Ishida et al., 2014; Pelosi et al., 2014; Iovinella et al., 2016). Other proteins commonly expressed in the chemosensory system include sensory neuron membrane proteins (SNMPs), which are related to scavenger proteins of the CD36 family (Benton et al., 2007), and several classes of G protein-coupled receptors (GPCRs) which allow neurons to sense a variety of extracellular signals, including e.g. hormones and neurotransmitters (Pándy-Szekeres et al., 2018).

Reverse genetic tools such as RNA interference (RNAi) has been widely used to validate the function of chemosensory genes (Palevich et al., 2018), and will be important to establish in the context of PRM. In PRM, RNAi silencing have previously been used to target gut associated genes for vaccine development (Kamau et al., 2013; Campbell, et al., 2012). However, there are no reports of RNAi targeting chemosensory tissues in any mite species including in this species. In this study, we make use of a non-invasive immersion-based methodology for achieving RNAi, and to show proof of concept that it is feasible to knock down the expression of a chemoreceptor transcript in *D. gallinae* olfactory neurons.

## 2. Material and methods

### 2.1 Mites

*D. Gallinae*, the poultry red mite, was a kind gift from Professor Baoliang Pan, China Agricultural University, Beijing, China. Upon arrival, mites were initially maintained at 24±2°C and 65%±4% relative humidity for 24 h, and then kept *in vivo* using a rearing system by feeding on chickens, as previously described (Wang et al., 2018). The care and use of chickens in this study was approved by Hainan University Institutional Animal Care and Use Committee.

### 2.2 Scanning electron microscope (SEM)

Adult mites and all pairs of legs were used for scanning electron microscopic examination, as previously described (Gainett et al., 2017). Briefly, specimens were cleansed with ultrasonic cleaning solution (Liquinox^®^) using an ultrasonic cleaner (Digital ultrasonic cleaner^®^). Samples were subjected to supercritical drying (Quorum^®^ K850) with CO_2_ after being dehydrated in a graded series of ethanol (from 45% to 100%). Dehydrated mites were fixed on stubs with double sided conductive carbon adhesive tabs and then sputter-coated with gold using Anhui Beq SBC-12 coating apparatus. Morphological analyses and photographs were done with a ZEISS GeminiSEM (field emission scanning electron microscope) at the Chinese academy of tropical agricultural sciences, Hainan, China.

### 2.3 Total RNA isolation, cDNA library preparation, and high-throughput sequencing

Total RNA extracted from the forelegs and hindlegs (forelegs consider as a treated group and hindlegs as a control group) of *D. gallinae mites* were used for RNA sequencing. A total of 150 adult mites (300 forelegs and 300 hindlegs) were dissected by using a scalpel blade (size no. 10) under a dissecting microscope, and three biological replicates per group were generated. After dissections, all samples were rapidly frozen using liquid nitrogen and kept at −80°C until use. RNA was extracted from each group using Trizol^®^ reagent (Thermo-Scientific, Carlsbad, CA, United States), according to the manufacturer’s protocol. The RNA quality was analyzed using an Agilent 2100 Bioanalyzer (Agilent tech., CA, United States), and high-quality samples were subsequently sequenced at the Beijing Genome Institute (BGI-Shenzhen, China), using a BGISEQ-500 platform. For generating first-strand cDNA libraries, RNA was purified using DNA polymerase I and deoxyribonucleotide triphosphate (dNTPs), following RNase H treatment. Then, the cDNA was generated using superscript III reverse transcriptase with random hexamer primer sets. Following sequencing, clean data (clean reads) were obtained by removing those containing adapter or unknown nucleotides at greater than 5% and of low-quality reads from the dataset. All clean reads were converted to the FASTQ file format. Simultaneously, the quality score (Q20 and Q30), GC contents and sequencing coverages were calculated. All of the downstream analyses were based on clean data with high-quality reads.

### 2.4 Bioinformatics and differentially expressed genes (DEGs) analysis

The reference genome of *D. gallinae* was obtained from NCBI (GAIF00000000) (Schicht et al, 2014) and used for building the index by HISAT2 (v2.0.4). Hierarchical indexing for spliced alignment of transcripts (HISAT) was based on the burrows-wheeler transform and the Ferragina-Manzini (FM) indices, using both genome-wide and local genome index forms (Kim et al., 2015). The generated index files were used to align the clean reads of the six RNA-seq samples to the reference genome. Bowtie2 (v2.2.5) was used to compare clean reads to reference sequences, and RSEM (v1.2.8) was used to calculate gene expression levels for each sample (Li and Dewey, 2011; Langmead et al., 2012). Gene expression levels were estimated based on the FPKM value (fragments per kilobase of exon per million fragments mapped) using RSEM (RNA-Seq by expectation-maximization) method with default parameters. Differential expression analysis of chemosensory genes in forelegs versus hindlegs were performed by the DEseq2 R package (Wang et al., 2010). The p-values were adjusted using q-values, with q-value ≤ 0.001 and log_2_ (fold change) ≥2 set as the threshold for significantly differential expression using the Benjamini and Hochberg method (Guo et al., 2008; Wang et al., 2010). Differentially expressed genes (DEGs) were cut-off by a false discovery rate (FDR) at 0.05, and then a gene ontology (GO) term enrichments were performed using Blast2GO software (Conesa et al., 2005). To clearly describe the DEGs of each olfactory-related family, we performed cluster analysis using pheatmap package in R software based on FPKM-values, represented as a heatmap.

### 2.5 Identification and annotation of chemosensory-related genes

Chemosensory-related genes from the transcriptomes of *D. gallinae* were identified by the functional annotation results based on its molecular function, cellular component and biological process, allowing for meta-analyses of gene populations. Standalone tblastn searches (BioEdit software 5.0.6) were performed using previously identified olfactory sequences of IRs, GRs, GPCRs, OBPs, NPC2 and SNMPs from other mite and tick species as queries in order to identify additional candidate chemosensory gene families in the available transcriptomes of *D. gallinae* (Hall, 1999; Ashburner et al., 2000). All candidate gene families were manually verified using the BLASTx (translated nucleotide to protein) program against non-redundant protein databases available at the NCBI with a cut-off e-value < 10^−6^. The open reading frames (ORFs) of all candidate genes were determined by using the open reading finder at NCBI. Thereafter, all transcripts with BLAST hits were included in an InterProScan tool search available at the EMBL-EBI with the following integrated database: ProfileScan, SignalPHMM, Phobius, HAMAP, FPrintScan, PatternScan, HMMPIR, Pfam, HMMPanther, HMMSmart, SuperFamily, patternScan, SFLD, TMHMM, Coils and CDD (Götz et al., 2008). The presence of putative N-terminal signal peptides were detected using the default parameter of SignalP4.0, while transmembrane domains were detected using the default parameter of TMHMM2.0. In addition, olfactory soluble proteins, such as OBP and NPC2 protein sequences from the arthropod that include insects, mites, ticks were used for motif-pattern analysis. The parameters setting in this study for motif predictions were: minimum width of motif = 6, maximum width = 10 and maximum number of motif to find = 8. Motif-based sequence analyses were predicted by using web-based version 5.1.0 of the MEME server (Machanick and Bailey, 2011). All candidate chemosensory unigene names were represented as follows: first letter of the genus and species (e.g. *Dermanyssus gallinae-Dg*) followed by a member of one of these protein families (e.g. IR, NPC2, OBP).

### 2.6 Phylogenetic analysis of chemosensory genes

For the qualitative report of gene family transcripts, the translated ORFs of *D. gallinae* chemosensory transcripts that contain the relevant conserved domains were retrieved and used to construct phylogenetic analysis. Amino acid sequences for each gene family were aligned using MAFFT in “Auto” strategy method with BLOSUM62 scoring matrix and other default parameters (Katoh et al., 2002). The maximum-likelihood (ML) trees were constructed with PhyML using the best-fit substitution model WAG+G+I as determined by ProtTest 2.4. Branch support was estimated using a fast and accurate maximum likelihood-ratio test (aLRT) and subsequently edited in iTOL webserver (Anisimova and Gascuel, 2006; Letunic and Bork 2007; Guindon et al., 2010). The species maximum-likelihood phylogenetic tree was conducted based on the alignment result of mitochondrial cytochrome c oxidase genes (CO1) from different species using the software in MEGA7.0 (Kumar et al., 2016). The tree image was further viewed and graphically edited using the iTOL online tool with the number of identified chemosensory genes.

### 2.7 Quantitative real-time PCR validation for DEGs

A total of 4 chemosensory-related transcripts identified by RNA-seq to be expressed deferentially between forelegs and hindlegs were selected for real-time quantitative PCR (qPCR) analysis using the LightCycler^®^ (Roche Applied Sciences, Mannheim, Germany). RNA extraction was performed as mentioned above. First-strand cDNA was synthesized with a PrimeScript™ RT synthesis kit with one-step genomic gDNA eraser kit according to instructions (TaKaRa, Tokyo, Japan). Gene-specific primer sets were designed based on the ORF, and Fun14-like gene was used as housekeeping gene (Bartley et al., 2012). Amplification was performed in a final reaction of 50 μl containing 5 μl of sample cDNA, 3 μl of each specific primer (10 mM), 25 μl of 2× SYBR green master mix (TransGen Biotech, Beijing, China), and sterilized ultra-pure grade ddH2O (14 μl). The thermal cycle conditions were as follows: 96 °C for 8 min, followed by 40 cycles of 96 °C for 5 s, 56 °C for 15 s and 71 °C for 15 s. Following the real-time PCR cycles, the specificity of the SYBR green PCR signal was confirmed by melting curve analysis and agarose gel electrophoresis. Relative quantification data were analyzed using the comparative 2^−ΔΔCt^ method with three biological repeats each containing three technical replicates (Livak and Schmittgen, 2001).

### 2.8 Preparation of double stranded RNA (dsRNA)

Total RNA was isolated from ten individual mites using Trizol^®^ reagent and was then treated with a PrimeScript™ RT synthesis kit with genomic gDNA eraser according to instructions (TaKaRa, Beijing, China). A 414 base pair (bp) fragment of the PRM OBP-like gene was amplified from the cDNA using the following the primer sets: (F-5’-ATAAGAAT<u>GCGGCCGC</u>ATGAAGGCAATCGTGCTTGTG-3’) and (R-5’-CCG<u>CTCGAGGGC</u>TTACTGTTCCACCTTGCACT-3’) containing EcoRI and XhoI restriction enzyme sites (Thermo-Scientific, Carlsbad, CA, United States). Two control groups were used: blank control (without any dsRNA) and negative control (mites exposed to a non-related dsRNA). The *β-glucuronidase* (GUS) gene (KY848224) was used as negative control group, as reported (Whyard et al., 2015). After purification of PCR products, the amplified PCR products were cloned into a T/A cloning vector PMD18-T (TaKaRa, Tokyo, Japan), and later excised from PMD18-T plasmid using NotI and XhoI restriction sites, then ligated into a similarly-digested plasmid pL4440, a vector possessing convergent T7 promoters. Positive colonies were transformed into *Escherichia coli* HT115 (DE3) competent cells, which are RNase-III-deficient strain, and then a single bacterial colony was cultivated in liquid LB medium containing 120 μgml^−1^ ampicilin and 13.5 μgml^−1^ tetracyclin for 12-15 hours at 37° C at 110 rpm. The culture was then diluted 50× in 1× LB broth and grown until OD_600_ value of 0.4-0.6. Isopropyl *β*-D-1-thiogalactopyranoside (IPTG) was added in a final concentration of 0.4 mM to induce T7 polymerase activity. The expressed dsRNA was extracted using Trizol reagent, and the quality of dsRNA were analyzed by electrophoresis on 2% agarose gel. The RNA concentration was quantified using a spectrophotometer (Implen NanoPhotometer N50). dsRNA aliquots were stored at −80 °C until used.

### 2.9 Delivery of dsRNA by non-invasive immersion method

Ten *D. gallinae* adult mites were immersed in separate sterile Eppendorf tube in 35-40 μl dsRNA encoding either DgOBP-1 (target gene) or GUS (negative control), diluted in saline water (0.9% Nacl) to a final concentration of 3.5 μg/μl, with 3 biological replicates per treatment and control group, as reported previously (Marr et al., 2015). The blank control group was directly immersed with DEPC-treated water. The treatment and control group were then incubated at 4 °C for 14 hours.

### 2.10 Validation of knockdown by real-time quantitative PCR

Real-time quantitative reverse transcription (qRT-PCR) assay was performed in triplicates. After 14 h of immersion, RNA extraction and first-strand cDNA synthesis were performed as described above. The cDNA from each replicate treatment was then used to assess the extent of RNAi by measuring levels of gene expression using qPCR. For relative quantification of expression, the Fun14-like gene was used as a housekeeping gene to compare levels of RNAi as described previously. The primer sequences were as follows: Fun14-like (F-5’-TAA CTG GCT CAC CCG AAC TC-3’ and R-5’-CTT TCT CCA CAG CCT TCC AG-3’; product size 208 bp) (Bartley et al., 2012). Melt curve analysis was performed and confirmed that only a single product was amplified with each primer pair in every sample. The 2^−ΔΔCt^ method was used to analyze gene expression profiles, comparing expression in specific dsRNA treated samples to *gus*-dsRNA treated samples (Livak and Schmittgen, 2001)

### 2.11 Data analysis

The data were analyzed using prism 5.01 software (GraphPad Prism, California, USA) and plotted as bar graphs, and the values are represented as means ± standard errors of means (SEM) of three biological replicates. The differences between two groups were analyzed using t-tests, and the differences between multiple groups were analyzed using one-way ANOVA followed by a Tukey’s multiple comparison test. Differences were considered to be significant at (*) p values of <0.05.

## 3. Results

### 3.1 Identification of sensilla on the legs of *D. gallinae*

Using scanning electron microscope, the forelegs and hindlegs of *D. gallinae* were analyzed, demonstrating a higher number of sensillum types on the anterior foreleg tarsi than on the posterior ones (Fig. 1B and C). Sensilla chaetica are classified into different morphological types based on location, size, shaft morphology, structure of socket and the presence and location of pores: sensilla chaetica with or without a tip pore (Sc-tp) and sensilla chaetica with wall pores (Sc-wp). The Sc-tp, which was the longest sensillum type, displayed a regular distribution pattern on the tarsi (Fig. 2A). This sensillum type has a prominent socket and an articulating membrane (Fig. 2B), with the sensillum wall having continuous longitudinal ridges along the shaft, and the shaft being gradually tapered toward the tip (Fig. 2D and E). Even though tip pores on the Sc-tp of the distalmost tarsomeres (DT) I-II were indistinct in the SEM (Fig. 2E), tip pores on DT III-IV were delimited by a round margin (Fig. 2F). In contrast to Sc-tp, all (Sc-wp) were restricted to tarsi I (Fig. 3A), and their shafts were inserted in a shallow membranous cuticle with an incomplete articulation (non-socketed) (Fig. 3B). The shaft tapered toward the tip (Fig. 3C), and the thick wall contained a large number of evenly distributed pores (Fig. 3E).

**Fig. 1.**
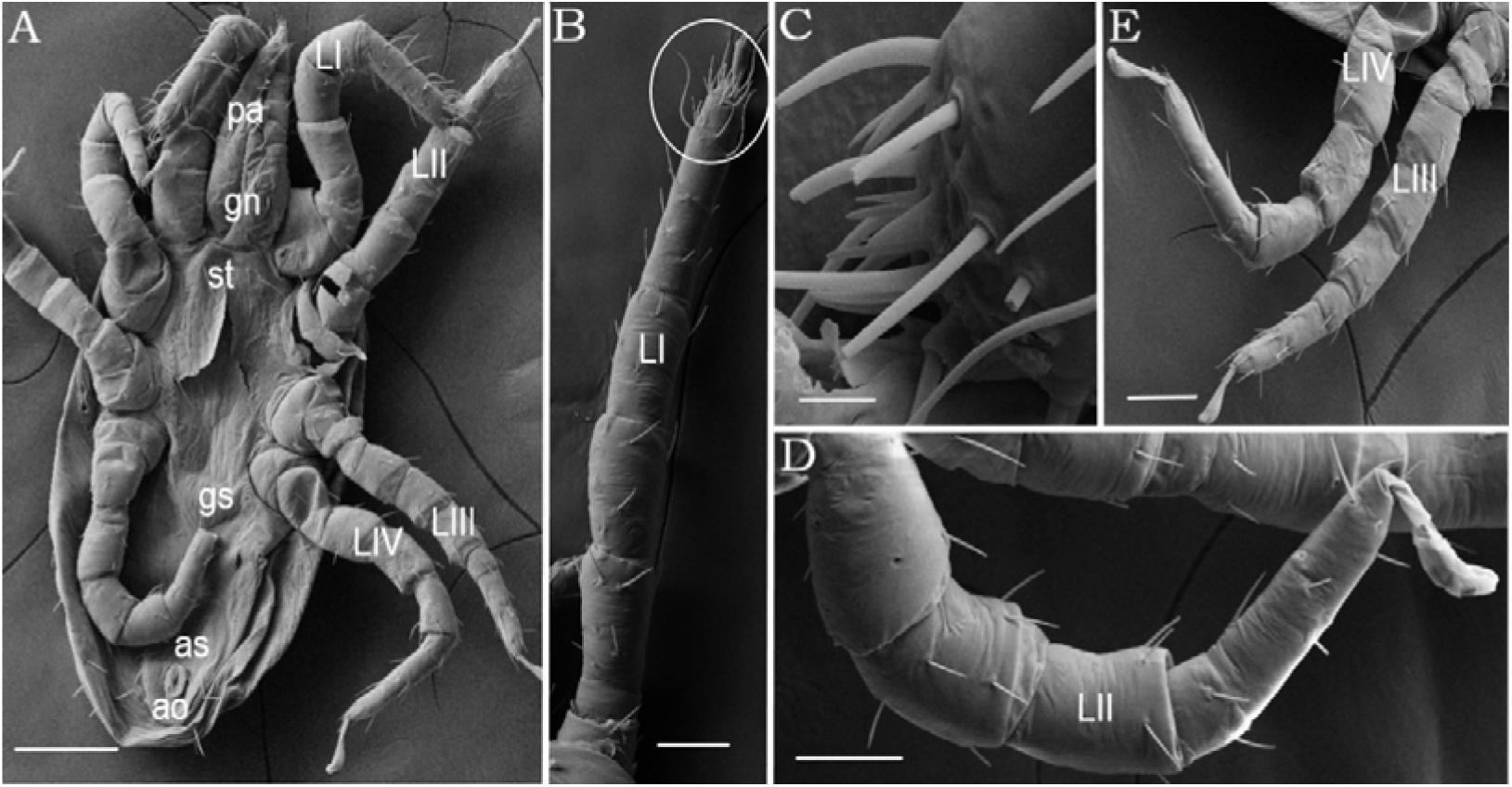
Scanning electron micrographs of PRM. A Ventral view of an adult PRM. B The first leg (LI), indicating the distal sensilla-rich structure (circle). C Different types of sensilla on the distal-most tarsomere (DT-I) of LI. D The second leg (LII). E The third leg (LIII) and fourth leg (LIV). Different types of sensilla are not discriminated in this figure because of low magnifications. *Scale-bars:* A, 100 μm B, D, E, F, 20 μm, C, 2 μm. *Abbreviations:* st, sternal plate; gn, gnathostoma; as, anal shield; ao, anal opening; ch, chelicera; Pa, pedipalp; (LI–IV), leg; gs, genitoventral shield

**Fig. 2.**
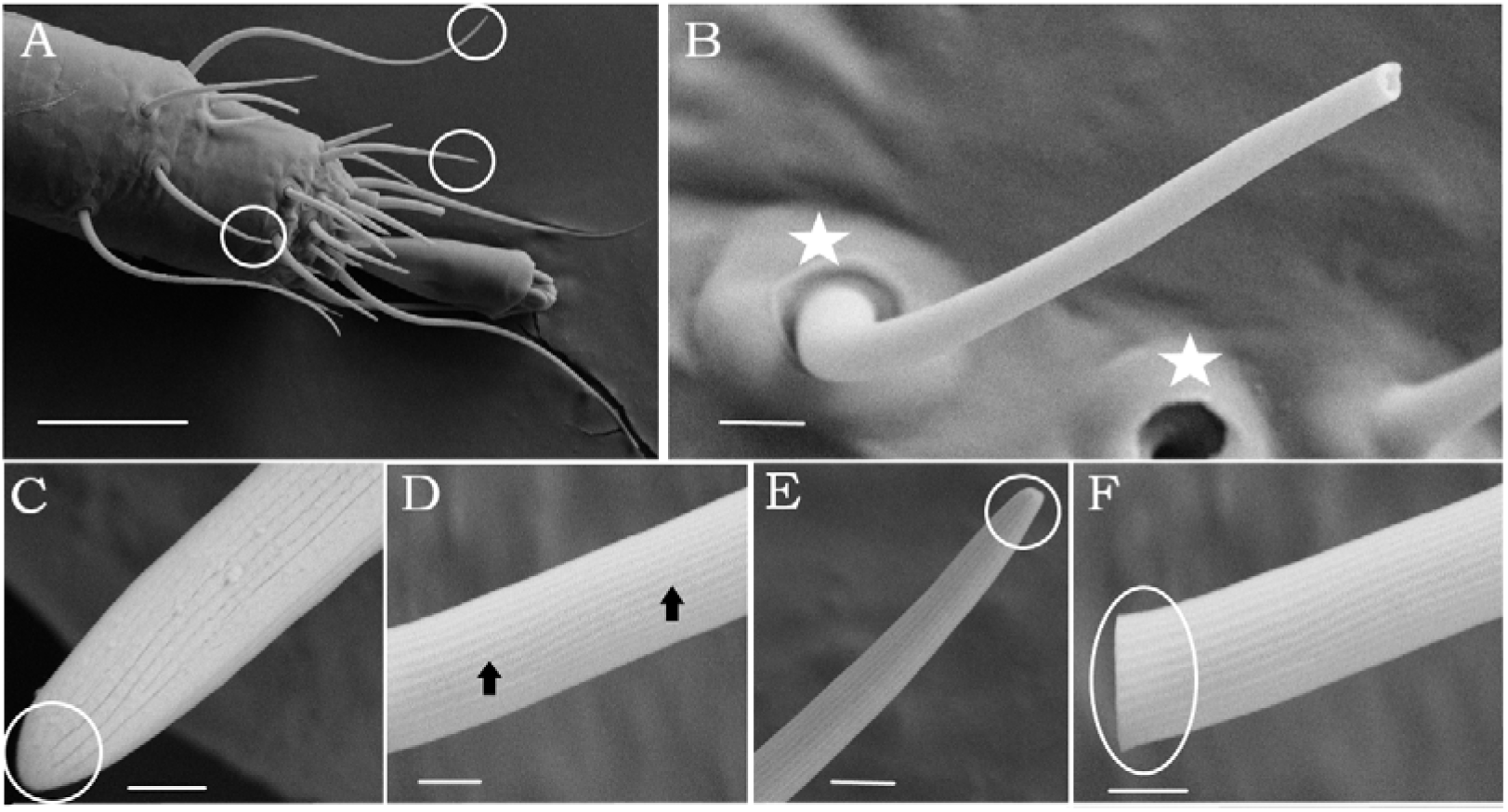
SEM of sensilla chaetica with or without tip-pore (Sc-tp). A. Circles indicating sensilla chaetica with a tip pore or without pores on distalmost tarsomere of first leg. Different types of sensilla are not discriminated in this figure because of low magnifications. B. Shaft broken in its apical part revealing a pore and structure of socket (□) C. no pore sensilla. D. Lateral view of the Sc-tp displaying longitudinal grooves (arrows). E. Tip-pore sensilla showed indistinct pore on tarsomere I (circle) F Tip-pore sensilla showed visible tip pore on tarsomere III and delimited by a round margin (circle). *Scale-bars:* A, 20 μm B, 2 μm, C, 300 nm, D,E,F, 1 μm.

**Fig. 3.**
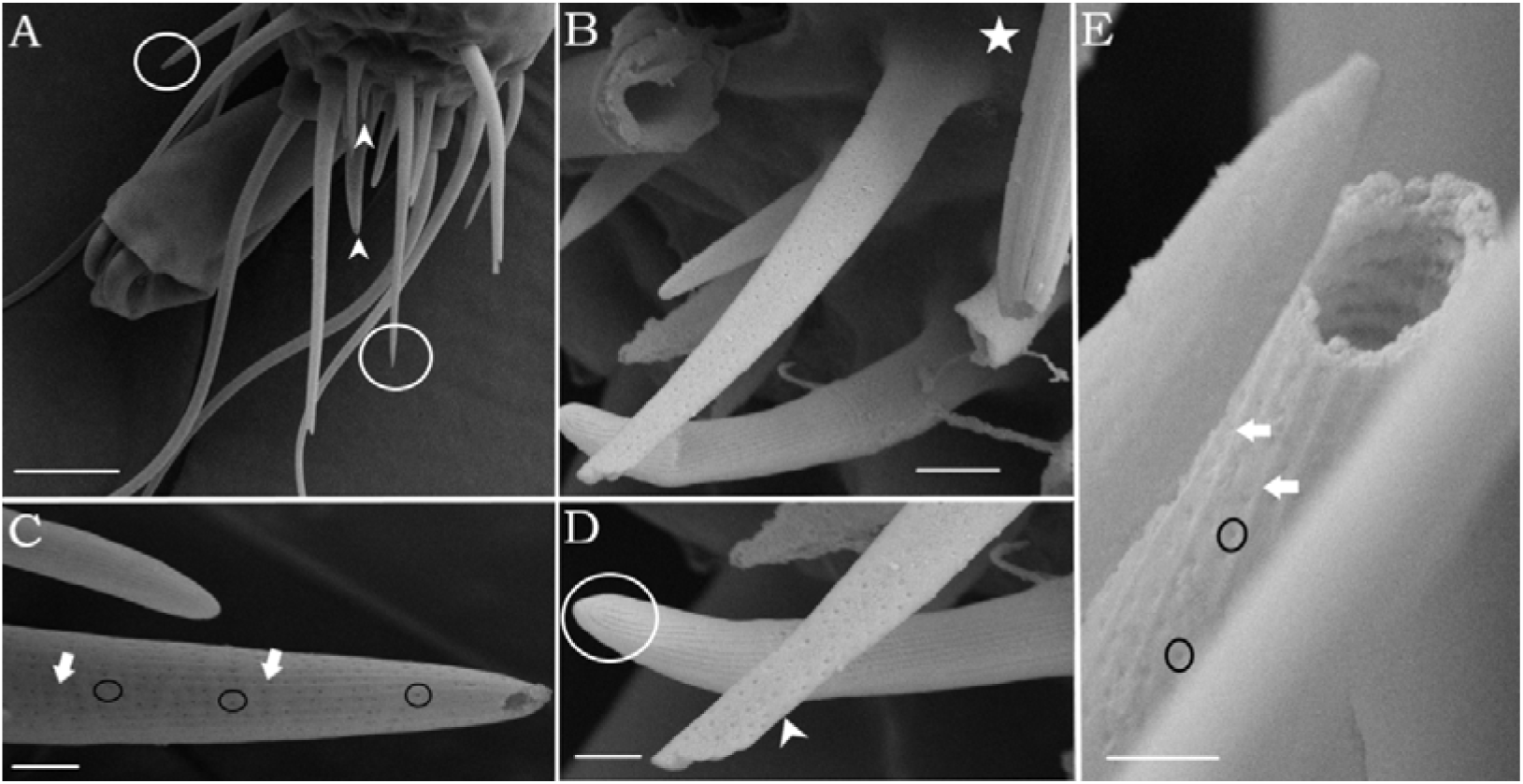
SEM of sensilla chaetica with wall-pores (Sc-wp). A. Lateral view of different types of hair sensilla on the distalmost tarsus (DT-I), circles indicate Sc-tp and arrowhead indicate Sc-wp sensillum B. Structure of socket with an incomplete articulation (□) C. shaft with longitudinal ridges (white arrows) and numerous pores housed in longitudinal grooves (black circle) D. comparing tip pore (circle) and wall pore sensilla (white arrows) E. White arrows represent longitudinal ridges, and circles indicate lines of wall pores. *Scale-bars:* A, 10 μm B, 2 μm, C, 1 μm, D, E, 200 nm.

### 3.2 Mapping and transcriptome analysis

To identify olfactory genes, RNA extracts of PRM forelegs and hindlegs were sequenced separately using a BGISEQ-500 HiSeq platform. On average, 22.5 and 21.7 million raw reads were generated for the foreleg and hindleg samples, respectively. After quality control screening, 69.42% of the reads per sample mapped to the *D. gallinae* reference genome (Table S1). While the resulting 25,604 sequences were identified as unigenes, these might not necessarily represent unique genes. The read sequences from the six transcriptomes were submitted to the sequence read archive (SRA) at NCBI under the accession number of PRJNA602095.

### 3.3 Homology and functional annotation

Of the 25,604 unigenes, 14,203 (55.47%) showed significant similarity to genes encoding for known proteins in the NCBI non-redundant protein database (NR), whereas 10,285 (40.16%) and 5,464 (21.34%) were identified following KEGG and GO annotation, respectively. According to the KEGG annotation results, signal transduction (environmental information processing) was the most highly expressed categories (Figure S1). GO annotation was used to categorize annotated genes into functional groups based on three main categories: molecular functions, cellular components and biological process. Among the biological process terms, the most represented biological processes were cellular (1,234 unigenes) and metabolic processes (9,54 unigenes). In the cellular component terms, the genes expressed were predominantly cell (1,309 unigenes), membrane (1,285 unigenes), and membrane part (1,221 unigenes). In the molecular function category, binding (1,477 unigenes) and catalytic activity (1,390 unigenes) had a huge preponderance and were the most highly expressed categories (Fig. 4). In general, GO term enrichment analysis indicated that forelegs were mostly associated with binding, catalytic activity, signal transducer and transporter activity in the molecular function category.

**Fig. 4.**
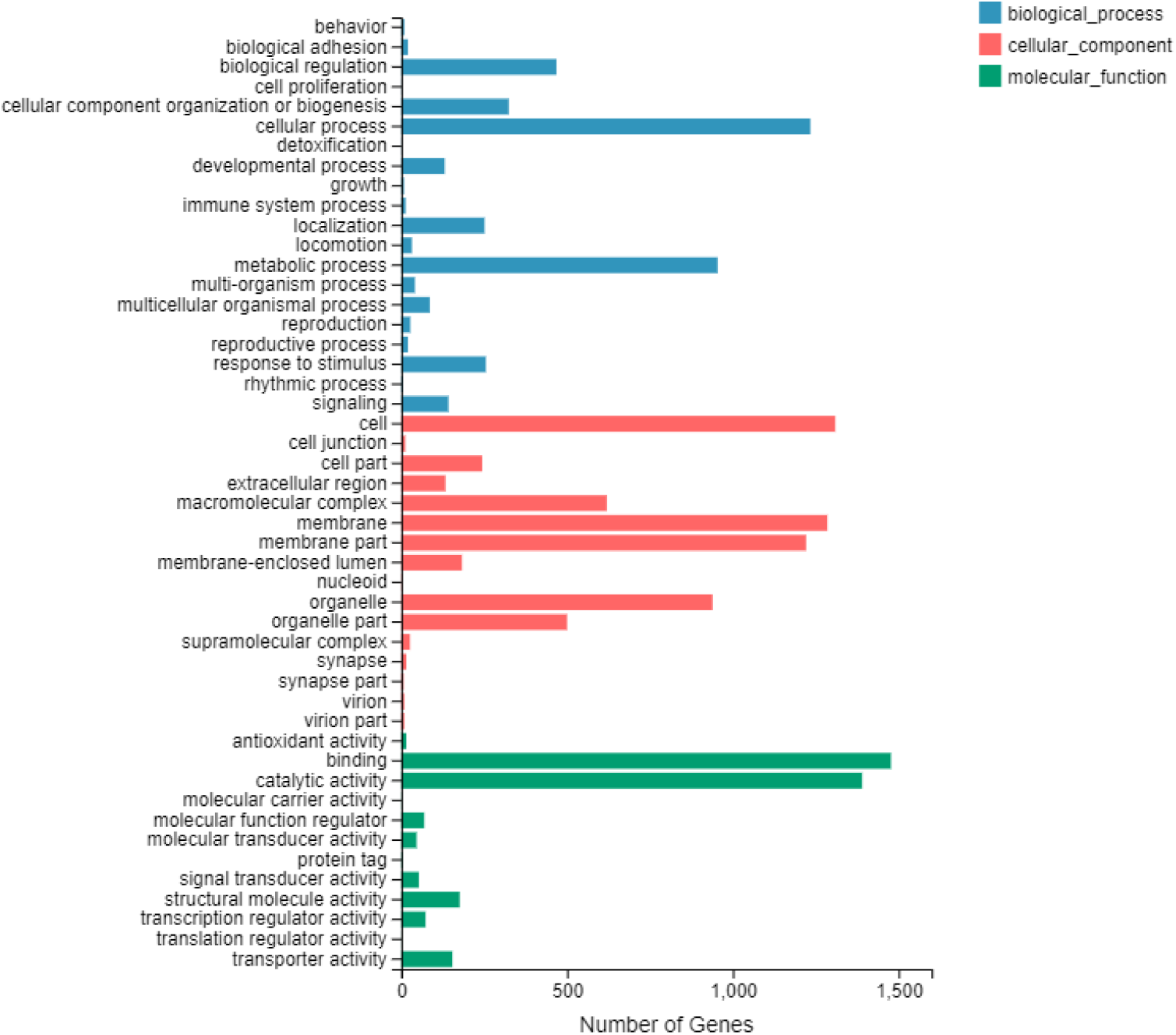
Gene ontology (GO) classifications of DEGs for forelegs vs hindlegs. X-axis represents the number of DEGs involved in particular GO term, and Y-axis indicates significantly enriched GO term.

### 3.4 No gustatory receptors (GRs), olfactory receptors (ORs) found in *D. gallinae* transcripts

BLASTx and BLASTn searches did not identify any transcripts putatively encoding for GRs and Ors in the *Dermanyssus* transcriptome. To further validate these findings, tBLASTn (*e*-value ≤ 1) and protein domain based searches did not identify any matches.

### 3.5 Identification of candidate IR family

Eleven transcripts encoding IR/iGluR homologs were identified, five of which contained the specific domain signature of the ionotropic glutamate receptors (Pfam domain-PF00060). Out of the eleven transcripts, only one transcript (DgIr3943) encoded all of the characteristic domains of the IR/iGluR proteins, namely, the amino-terminal domain (Pfam domain-PF01094), the ligand-binding domain (Pfam domain-PF10613) and the ion channel domain (Pfam domain-PF00060). The protein domain organization of iGluRs/IRs in PRM is shown in histogram (Fig. 5a). Among the IRs, all sequences showed complete ORFs, with >112 amino acids. However, read counts for the identified IR/iGluR transcripts were low in both forelegs and hindlegs: in the forelegs, IR transcript expression levels ranged from 0.06 to 0.13 FPKM, and in the hindlegs from 0.01 to 1.12 FPKM. None of the identified IR/iGluR transcripts were significantly found in the forelegs (Fig. 5B). To further distinguish putative IR gene families from the transcriptome of *D. gallinae*, all of the IRs in our transcriptomes were aligned with IRs from *Varroa destructor, Tetranychus urticae, Galendromus occidentalis*, *Ixodes scapularis, Tropilaelaps mercedesae* and *Drosophila melanogaster* by MAFFT (Table S2). The phylogenetic analysis demonstrated that eight genes were clustered with the iGluR sub-group, and one gene was close to the NMDA sub-group. Apart from that, two *Dermanyssus* transcripts formed a distinct sub-group together with a sequence of *Varroa* mites, suggesting that a mite-specific IR sub-group is present. The IR phylogeny did not reveal any potential orthologous relationships with insect IRs, for example there were no relatives of the divergent class of insect IRs (Fig. 6).

**Fig. 5.**
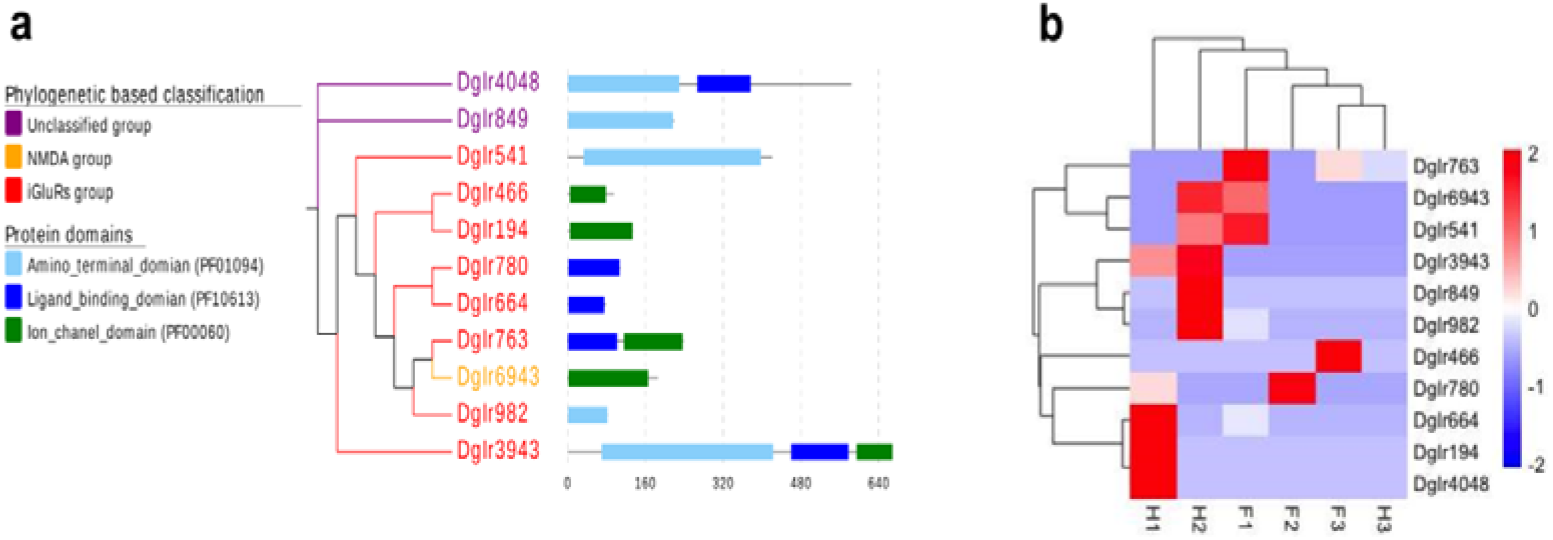
a The protein domain organization of iGluRs/IRs in PRM is shown in histogram. Phylogenetic tree of the IR/iGluR family based on PRM amino acid sequences, and the corresponding protein domains of each member. b Hierarchical clustering of the differentially expressed genes (DEGs). Blue to red colors represent gene expression levels (i.e., FPKM values from −2 to 2)

**Fig. 6.**
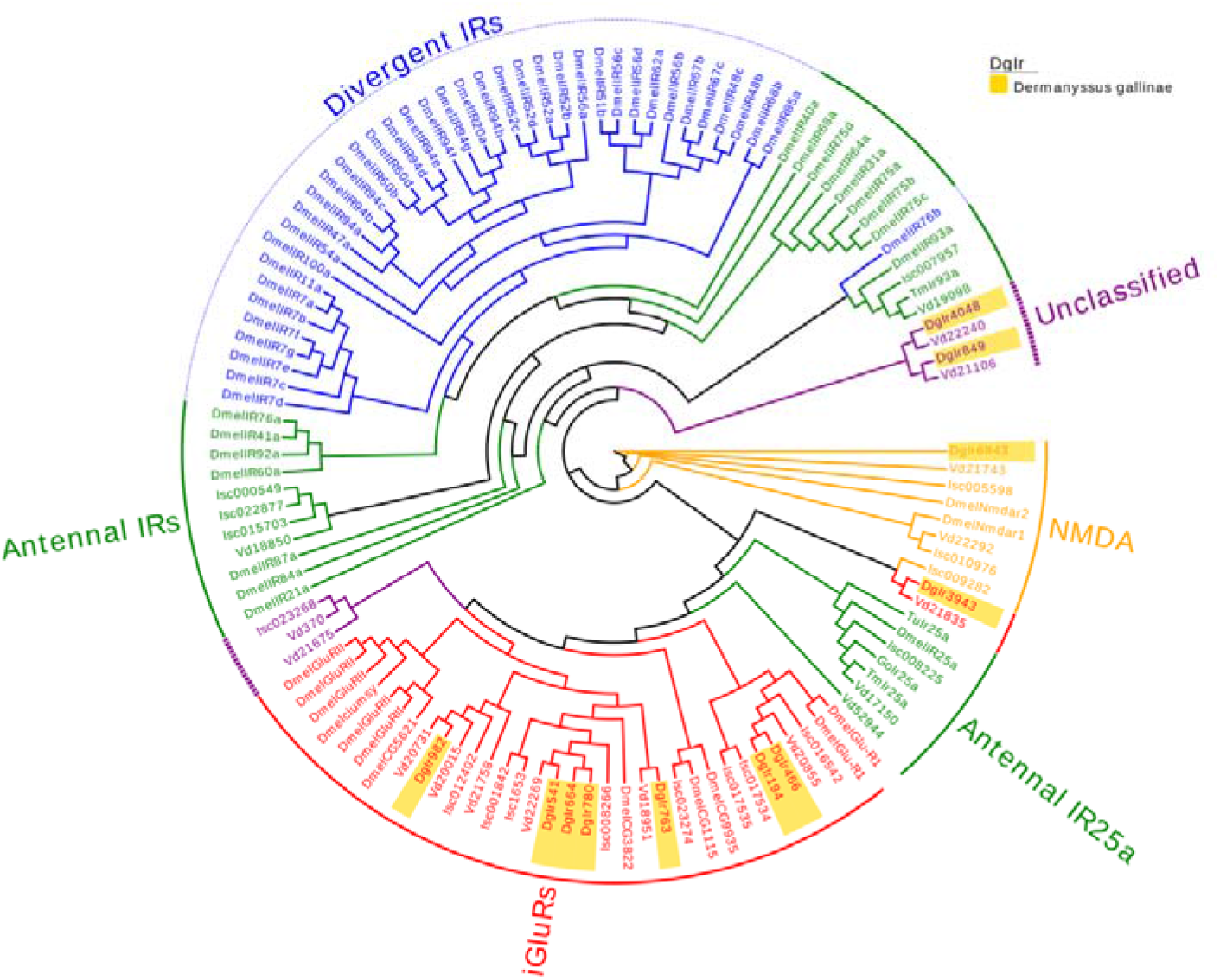
Phylogenetic tree of putative *D. gallinae* iGluRs and IRs along with insect and non-insect arthropods iGluRs and IRs. The *D. gallinae* genes are shown in light orange. Branch support was estimated by an approximate likelihood-ratio test (aLRT) (circles > 0.80). The amino acid sequences and a comprehensive list of acronyms are listed in S2.

### 3.6 Identification of putative OBP-like and NPC2 proteins in *Dermanyssus* sequences

A total of five and six candidate OBP- and NPC2-encoding genes were identified, respectively, in the transcriptomic analysis of *Dermanyssus* chemosensory tissues. These genes were similar to those reported for other mite species (Eliash et al., 2017). The complete ORFs were identified for five OBP candidate genes, revealing >137 amino acids encoding for proteins with high homology to known OBP-like proteins from *V. destructor* (XP_022653281.1; XP_022653293.1; XP_022645714.1; XP_022666940.1; XP_022672530.1), *Amblyomma americanum* (JZ172282.1) and *I. scapularis* (XP_002433530.1). SignalP revealed an 18-amino-acid-long signal peptide cleaved between position 18 and 19 (data not shown). Alignment of amino acid sequences from these species revealed that all OBP-like sequences contain six conserved cysteine (C) residues, with the exact same spacing pattern of all cysteines “C1-X16-C2-X18-C3-X37-C4-X15-C5-X9-C6” (where X represents any amino acid; prosite pattern notation, http://prosite.expasy.org/prosuser.html#conv_pa) (Fig. 7a).

**Fig. 7.**
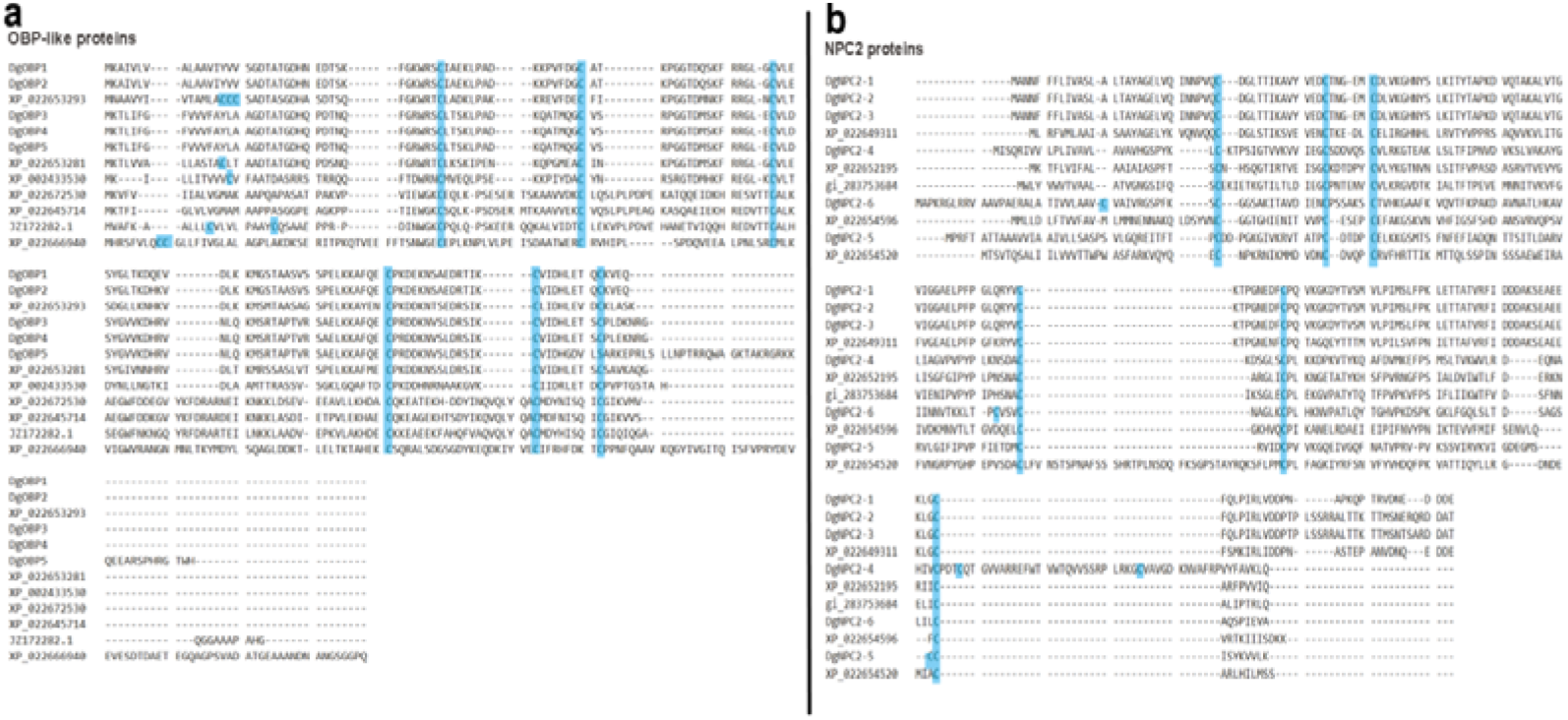
a Alignment of the five putative OBP-like sequences of PRM together with those of *V*. *destructor* sequences, *A. americanum* and *I. scapularis*. The highly conserved six cysteine (C) residues are shown in blue. b Alignment of the six putative NPC2 sequences of PRM together with those of *V. destructor* sequences. The highly conserved six cysteine (C) residues are shown in blue.

Two OBP-like genes (DgOBP-1 and DgOBP-2) revealed higher levels of expression in forelegs, whereas three genes (DgOBP-3, DgOBP-4 and DgOBP-5) had expression in both forelegs and hindlegs. Gene expression levels of all OBP-like proteins identified from the forelegs and hindlegs transcriptomes were compared (Fig. 8b). The phylogenetic analysis showed that all of *Dermanyssus* OBPs were clustered together on the same subclade with high bootstrap support. In addition, *Dermanyssus* OBPs were clustered with a clade containing OBPs of the mites *V. destructor* and *T. mercedesae*, with more than 80% branch support (Fig. 8a and Table S3). Conserved OBP motifs were predicted to better understand the protein’s evolution and function. Since a high number of OBP genes have been reported in both insects and arachnids, we performed a motif-pattern analysis between these two classes (Table S4). Only motif 1 (CMDYHJSQIC) and motif 2 (TCALKSEGWF) encoded the OBP family were present in all the OBP orthologs except for *Trichomalopsis sarcophagae* (Ts2), *D. willistoni* (Dw) and *D. ficusphila* (Df) (Fig. 9). Since motif 1 and 2 were found in all OBP proteins including the five *Dermanyssus* OBPs, the results provide confidence of their identification as bonafide OBP-like encoding genes and infer a functional similarity with other OBPs.

**Fig. 8.**
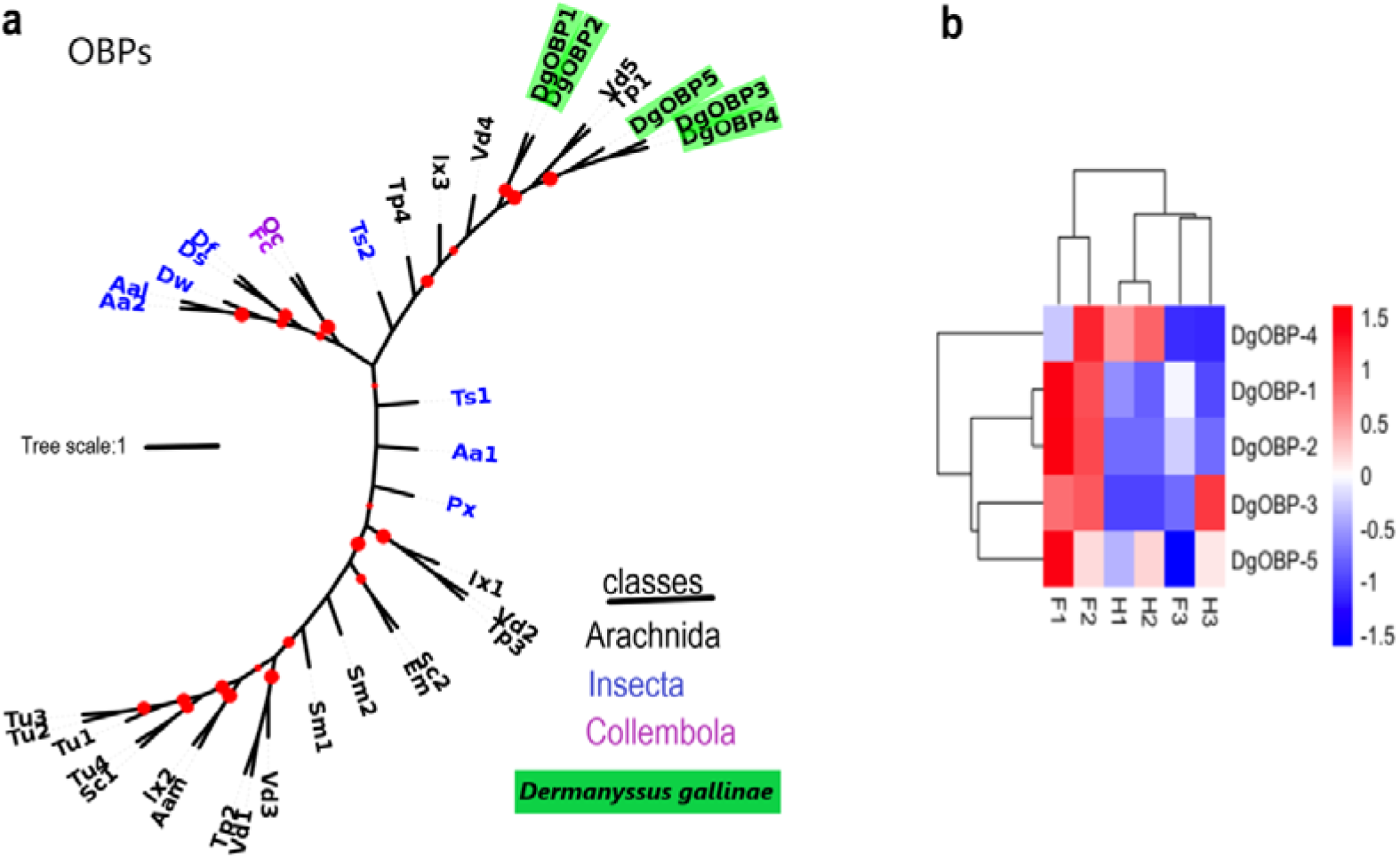
a Phylogenetic unrooted tree of odorant-binding protein (OBPs). Included are OBPs from *D. gallinae* (Dg: 5), *V. destructor* (Vd: 5), *I. scapularis* (Ix: 3), *A. americanum* (Aam: 1), *T. mercedesae* (Tp: 4), *Stegodyphus mimosarum* (Sm: 2), *Tetranychus urticae* (Tu: 4), *Sarcoptes scabiei* (Sc: 2), *Trichomalopsis sarcophagae* (Ts: 2), *Aedes albopictus* (Aal: 1), *Aedes aegypti* (Aa: 2), *Papilio xuthus* (Px: 1), *Folsomia candida* (Fc: 1), *D. sechellia* (Ds: 1), *D. ficusphila* (Df: 1), *Euroglyphus maynei* (Em: 1), *Orchesella cincta* (Oc: 1) and *D. willistoni* (Dw: 1). Bootstrap values were estimated using a recent fast approximate likelihood-ratio test (aLRT) and are displayed as circles on the branches, and are proportionate to the circle diameter (0–500). Leaves are coloured according to the organisms classes: arachnida (black), collembola (purple) and insecta (blue). *Dermanyssus* transcripts are highlighted in green. The amino acid sequences and a comprehensive list of acronyms are listed in S3. b Hierarchical clustering analysis of the differentially expressed genes (DEGs). Blue to red colors represent gene expression levels (i.e., FPKM values from −1.5 to 1.5).

**Fig. 9.**
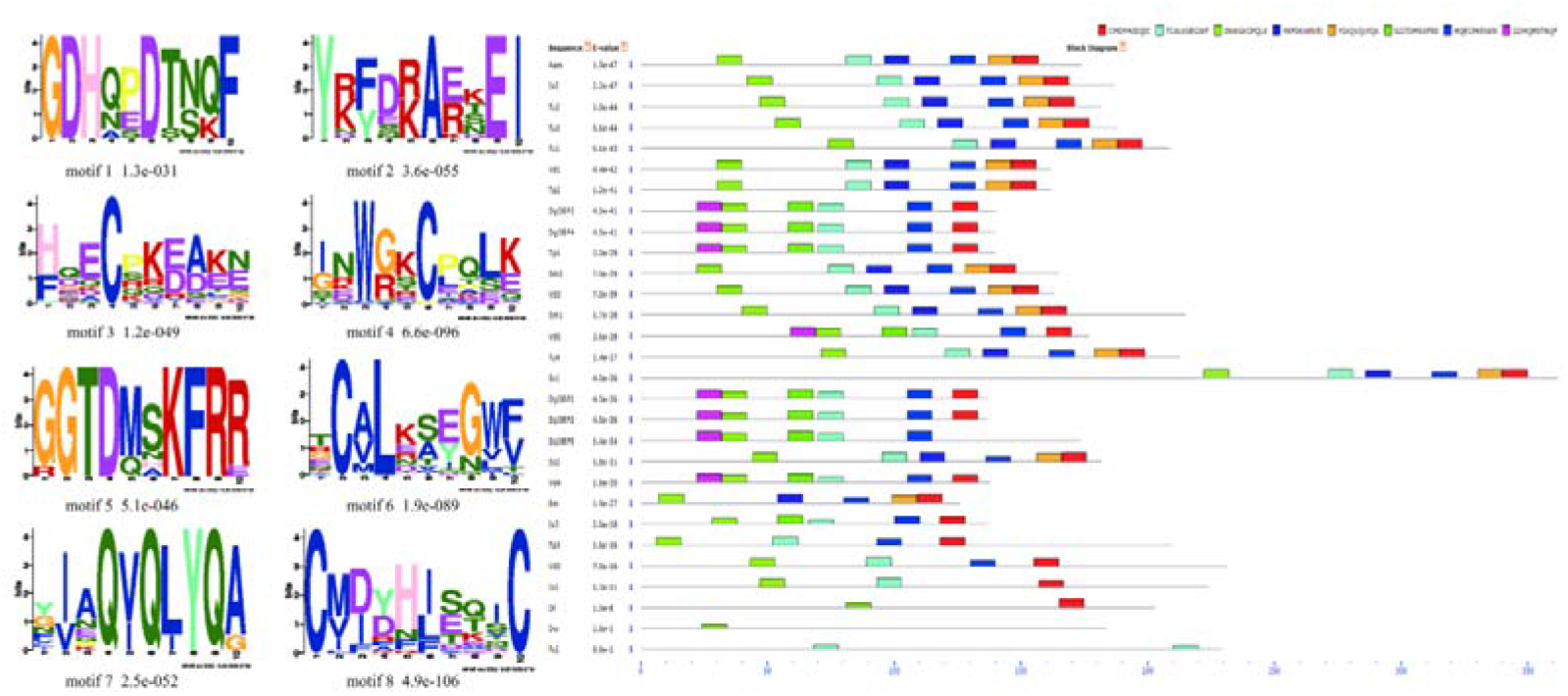
Motif-pattern analysis of OBP protein from *D. gallinae* and other classes including insecta and arachnida. Eight amino acid motifs with various widths and corresponding e-values were identified, and the lower part indicates approximate locations of each motif on the protein sequence. Different motifs are shown by different colored boxes, where small number indicates high conservation. Protein sequence of OBPs used in this analysis are listed in table S4.

Six NPC2 proteins were annotated from the *D. gallinae* transcriptomes. These putative NPC2 genes consisted of complete ORFs with lengths ranging from 471 bp to 591 bp nucleotides, and all sequences had an N-terminal signal peptide that are a common feature of the secretory proteins. To compare sequence similarity with other acari, a multiple sequence alignment showed that all six cysteines were conserved in all NPC2 proteins (Fig. 7b). Phylogenetic analysis of the NPC2 genes revealed three groups of *D. gallinae* NPC2 genes, with two groups being unique to arachnida (Fig. 10a and Table S5). The differential expression of the chemosensory-related proteins between the forelegs and hindlegs comparisons was less prevalent. Only a single NPC2 transcript, with a low abundance, demonstrated a foreleg-biased expression (Fig. 10b).

**Fig. 10.**
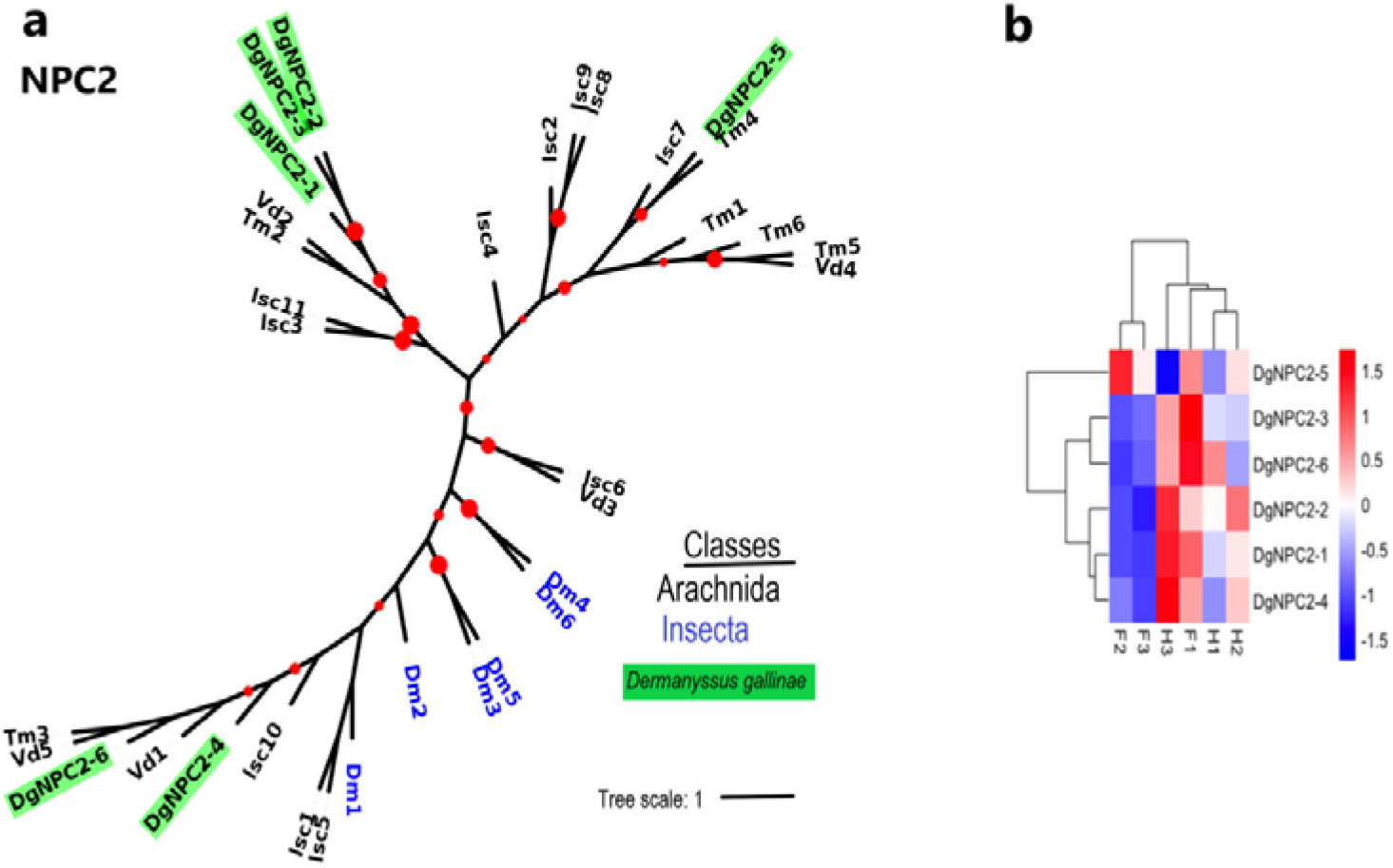
a An unrooted phylogenetic tree of NPC2 proteins. The evolutionary tree is constructed by aligning the SNMP sequences of *V. destructor* (Vd: 6), *Dermanyssus gallinae* (Dg: 6), *Drosophila melanogaster* (Dm: 6), *Ixodes scapularis* (Isc: 11) and *Tropilaelaps mercedesae* (Tm: 6). Bootstrap values were estimated using a recent fast approximate approximate likelihood ratio test (aLRT) and are displayed as circles on the branches, and are proportionate to the circle diameter (0–500). Leaves are colored according to the organisms classes: Insecta (blue) and Arachnida (black). *Dermanyssus* transcripts are highlighted in green. The amino acid sequences and a comprehensive list of acronyms are listed in S5. b Hierarchical clustering analysis of the differentially expressed genes (DEGs). Blue to red colors represent gene expression levels (i.e., FPKM values from −1.5 to 1.5).

The conserved motifs are important elements of functional domains. A total of 32 sequences from both insects and arachnids including six *Dermanyssus* NPC2 were used to compare and search for shared motif patterns (Table S6). The results showed that motif 1 (CPLKKGKDYT) and motif 2 (PFPGPKSDAC) were present in all the NPC2 orthologs except for *I. scapularis* (Isc1), *Tropilaelaps mercedesae* (Tm3,1), *D. melanogaster* (Dm2,4), *D. gallinae* (DgNPC2) and *V. destructor* (Vd4) (Fig.11). In addition, the homologous NPC2 from different species had similar motif patterns; for example, *D. gallinae* (DgNPC2-1, 2 and 3), *V. destructor* (vd2) and *T. mercedesae* (Tm2) had same motif-pattern as 4-7-2-8-1-6-3.

**Fig. 11.**
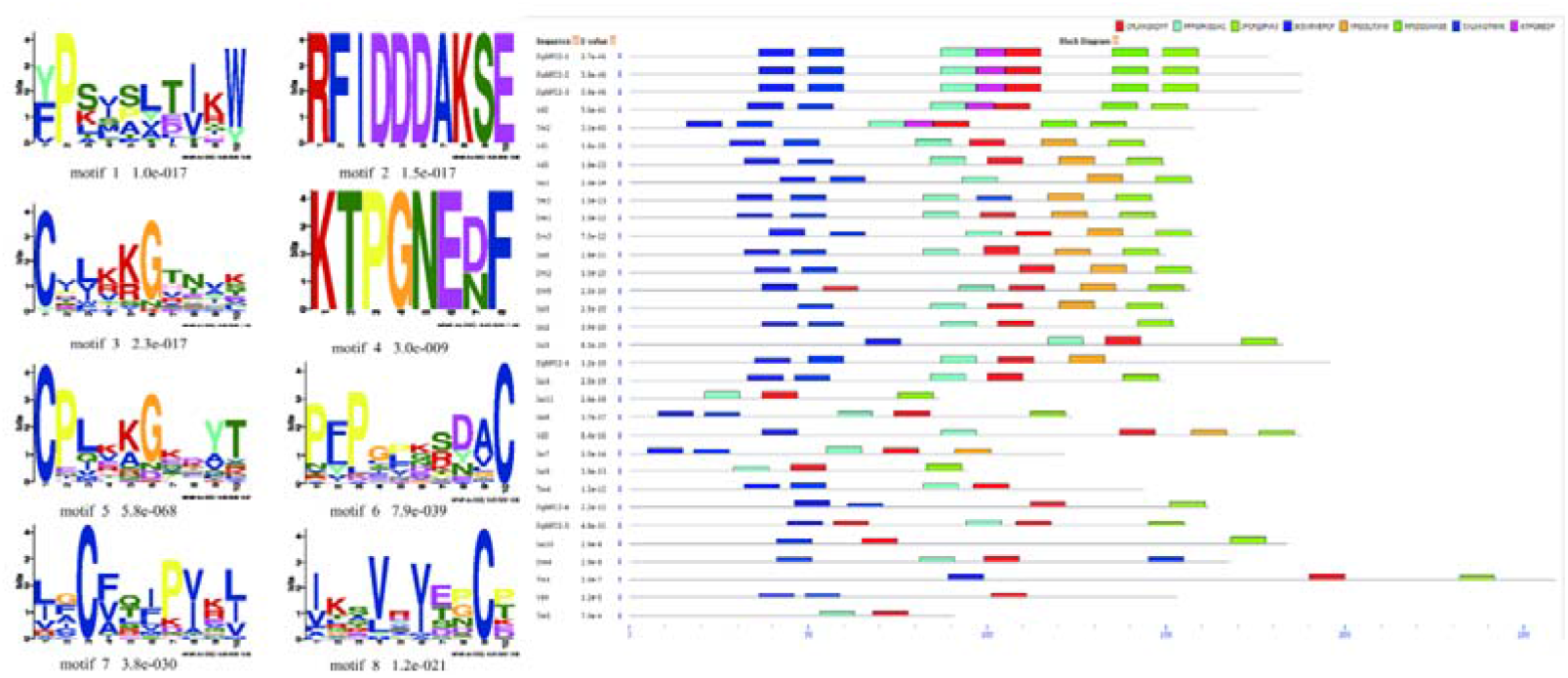
Motif analysis of NPC2 protein from *D. gallinae* and other classes including insecta and arachnida. Eight amino acid motifs with various widths and corresponding *e*-values were identified, and the lower part indicates approximate locations of each motif on the protein sequence. Different motifs are shown by different colored boxes, where small number indicates high conservation. Protein sequence of NPC2 used in this analysis are listed in table S6.

### 3.7 Identification of candidate sensory neuron membrane proteins (SNMPs)

Five candidate SNMPs transcripts were identified based on similarity to known SNMPs, within the CD36 superfamily (PF01130.21), of mites, ticks and insects. None of the identified transcripts were significantly found in the forelegs (Fig. 12b). To extend and specify this classification, the constructed phylogenetic tree revealed that acarian SNMP sequences were clearly separated from those of insects with the exception of three SNMPs (TrpWZC0, Var27012 and IscPD82) that showed proximity to the *D. melanogaster* clusters (Fig. 12a and Table S7).

**Fig. 12.**
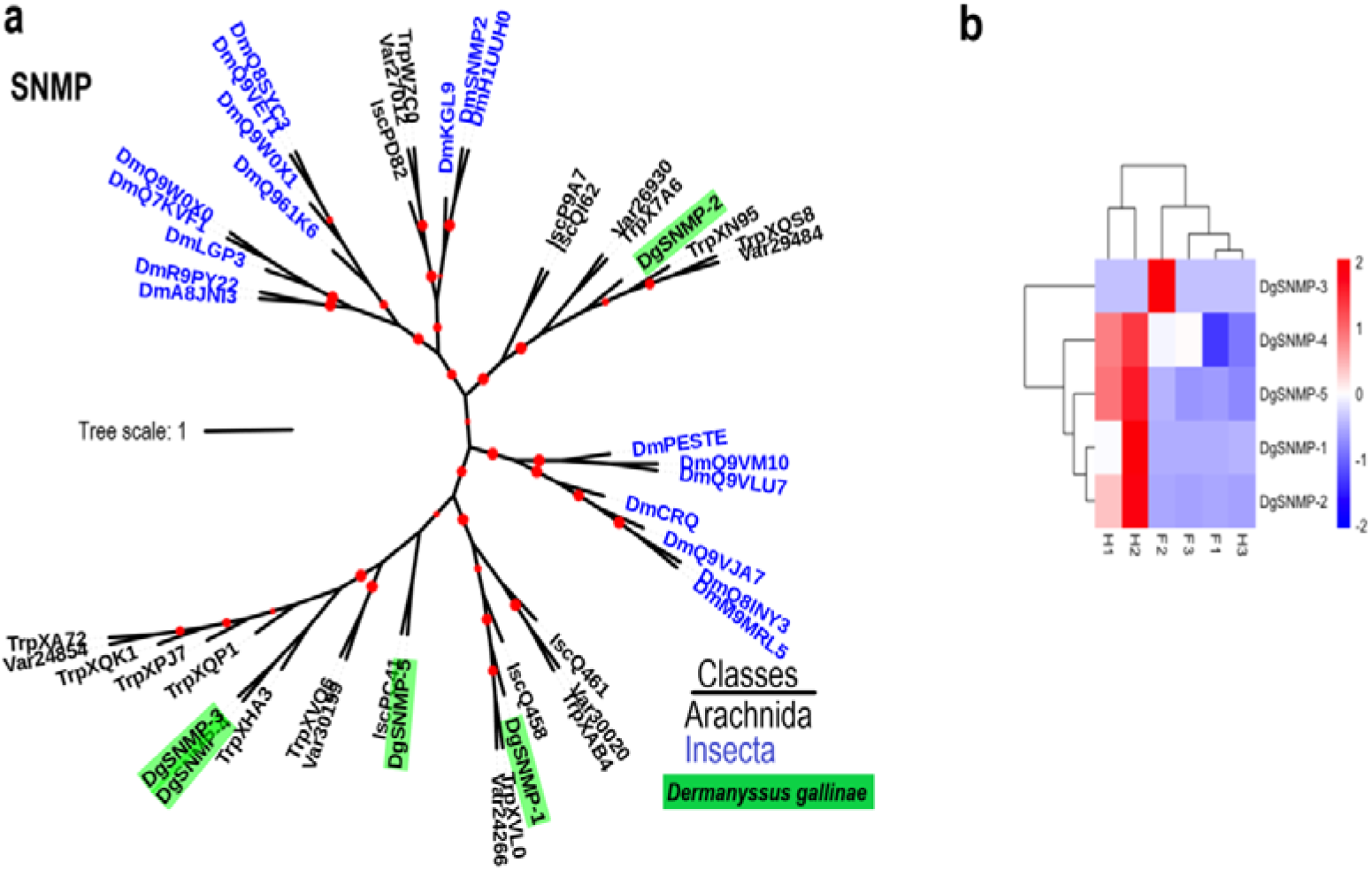
a An unrooted phylogenetic tree of SNMP proteins. The tree is constructed by aligning the SNMP sequences of *V. destructor* (Var: 7), *D. gallinae* (Dg: 5) *T. mercedesae* (Trp: 12), *Ixodes scapularis* (Isc: 6) and *Drosophila melanogaster* (Dm: 19). Bootstrap values were estimated using a recent fast approximate likelihood ratio test (aLRT) and are displayed as circles on the branches, and are proportionate to the circle diameter (0–500). Leaves are coloured according to the organisms classes: Insecta (blue) and Arachnida (black). *Dermanyssus* transcripts are highlighted in green. b Hierarchical clustering analysis of the differentially expressed genes (DEGs). Blue to red colors represent gene expression levels (i.e., FPKM values from −2 to 2).

### 3.8 Identification of candidate G protein-coupled receptors (GPCRs)

One putative GPCR was identified exclusively in the transcriptome of forelegs and contained a rhodopsin-like domain (PF00001.21). Protein domain and phylogenetic analyses of the putative GPCR transcript revealed that the transcript belongs to the putative clade A, rhodopsin-like GPCRs showing GPCR and photoreceptor activity. In insects, GPCRs are divided into three major classes, rhodopsin (clade A), secretin (clade B), metabotropic glutamate (clade C) and atypical (clade D) (Carr et al., 2017) (Fig. 13 and Table S8). We also found foreleg-biased expression of a G protein, and expression of this was verified by RT-PCR (qPCR) (Fig. 14).

**Fig. 13.**
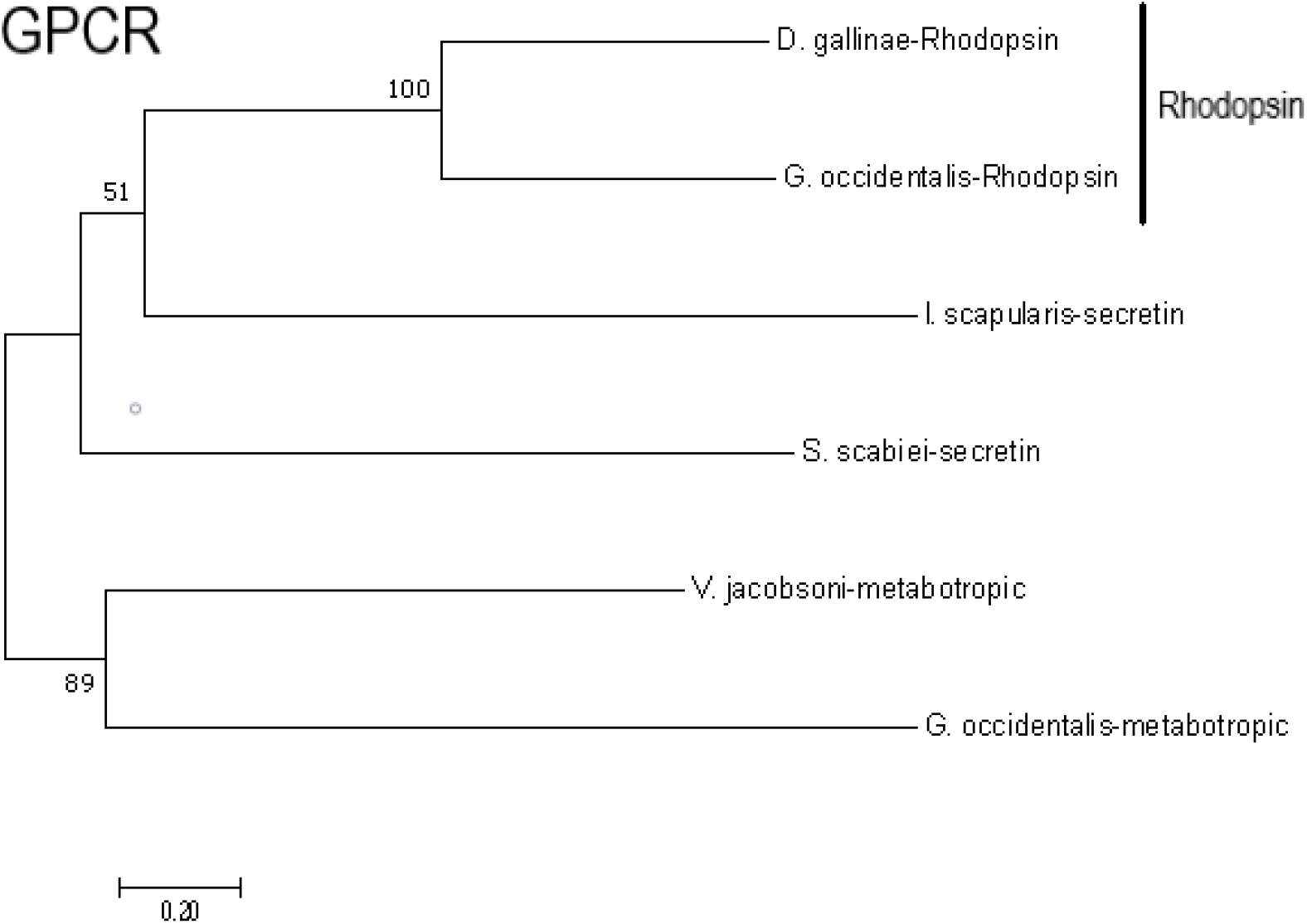
Phylogenetic tree of rhodopsin-like GPCR of *D. gallinae* and other mite species. The amino acid sequences are listed in S8.

**Fig. 14.**
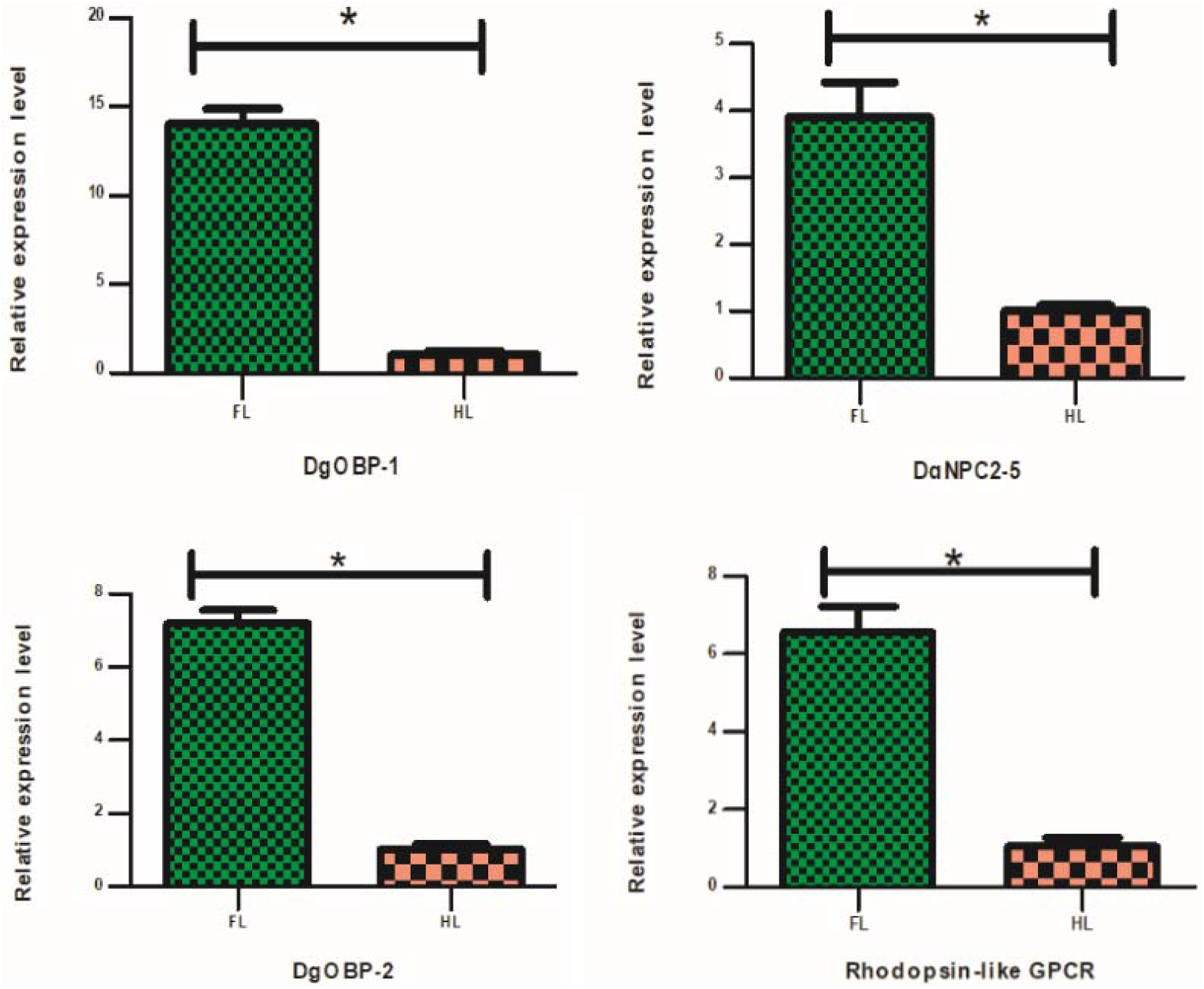
Expression profiles of putative chemosensory genes in *D. gallinae* using qRT-PCR. FL, forelegs; HL, hindlegs. Error bars represent means ±SEM, and asterisks above indicate significant differences between forelegs and hindlegs (*p < 0.05).

### 3.9 Differentially expressed gene (DEG) validation using qRT-PCR

In order to verify the differentially expressed genes identified through the transcriptome analysis, the expression pattern of identified gene were analyzed using qRT-PCR, including 2 OBPs (DgOBP-1, DgOBP-2), 1 NPC2 (DgNPC2-5) and 1 GPCR (DgGPCR-1). The expression levels were largely consistent with the transcriptome profile analyses concluding that the RNA-seq data were reliable (Fig. 14). The primers used for real-time PCR are listed in Table S9.

### 3.10 The efficiency of RNAi in PRM

DgOBP-1 was used as a test gene to assess the feasibility of using a dsRNAi gene knockdown approach to knock down gene expression in *D*. *gallinae*. As shown in Fig. 15, mites immersed in OBP dsRNA for 14 hours exhibited significant gene knockdown when compared to controls. Mites that were immersed in blank or negative control, exhibited no significant differences in OBP mRNA levels, as determined by qRT-PCR. The primers used for RNAi and real-time PCR are listed in Table S10.

**Fig. 15.**
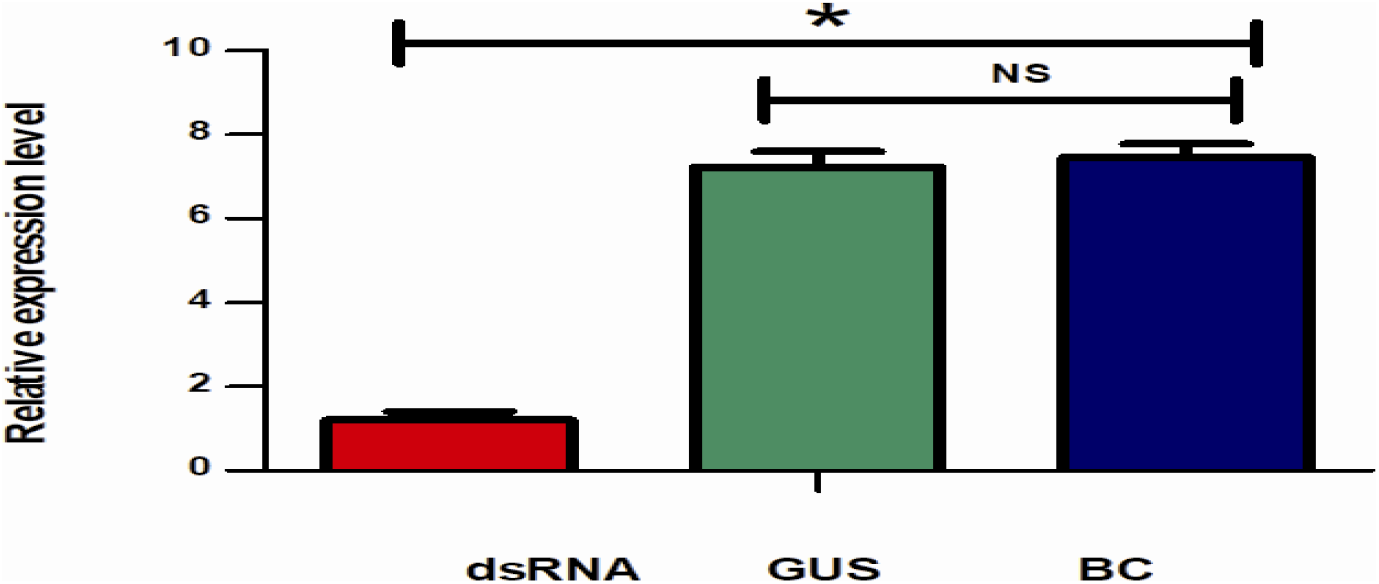
Immersion-based gene silencing of candidate DgOBP-1 in *D*. *gallinae* by RNAi. Results are presented as the mean ± the SEM. Statistical differences relative to the control group are presented as *p–value < 0.05; NS, no statistical significance. Abbreviations: dsRNA, treatment of mites with ds-OBP; GUS, treatment of mites with ds-GUS RNAs; BC, treatment of mites with DEPC-treated water (without any dsRNA).

## 4. Discussion

Compared to other arthropod in general and to ticks in particular, the molecular basis of chemoreception in mites is largely unexplored. Most arachnids including PRM are active at night and thus do not primarily depend on visual sense. Their most important sensory input comes from mechanical stimuli such as air currents, touch and vibrations. To better understand how mites perceive olfactory chemical cues, chemosensory genes were identified using transcriptomic analysis of *D*. *gallinae*, along with an ultrastructural study of tarsal sensilla. Our results suggest that the forelegs are equipped with sensilla containing neurons with chemo- and mechanosensory related functions. As a means to gain insight into the function of the chemosensory system of the PRM, an RNAi silencing approach was developed. There, we demonstrate a clear knock-down in the relative expression of the OBP-1 gene in dsRNA treated mites. In general, an improved understanding of the olfactory system may provide a good start of controlling this problematic mite.

Three types of setiform sensilla were distinguished, which is typical for all acari: no pore (np), wall-pored (wp) and tip-pored (tp) sensilla. Although each sensillum type has a characteristic ultrastructure, functional properties can only be established with certainty by data from physiological recordings, which we so far lack. Wall-pore sensilla are, in general, considered to have an olfactory function (Altner et al., 1977; Altner and Prillinger, 1980). The exclusive presence of wp sensilla on the distal-most tarsomeres of the front legs indicate that these are important sensory (Fig. 3) appendages also present in ticks (Haller’s organ) and other mite species (Eliash et al., 2017; Carr et al., 2017; Lei et al., 2019). Moreover, the presence of tp sensilla suggests a contact-chemoreceptive (gustatory function) and contact-mechanoreceptive function (Fig. 2), as previously described in other arachnids. In contrast to wp and tp sensilla, np sensilla are considered to be mechanosensory function to sense a mechanical distortion of the exoskeleton or seta (Fig. 2C) (Altner and Prillinger, 1980). These results are consistent with ticks and other types of arachnids (Gainett et al., 2017; Lei et al., 2019).

In forelegs, four molecular function categories might be connected to olfaction (odorant binding), enzyme activity (catalytic) and signal transduction (signal transducer activity) (Fig. 4). These results are likely to reflect that forelegs play an important chemosensory role. Nevertheless, GO annotations are common to all tissues types that do not provide any information on the expression levels of the individual gene types. Our study did not identify any transcripts putatively encoding gustatory receptors, despite well-documented morphological evidence of gustatory-like tp sensilla (Fig. 2e). This result might reflect that the transcriptome sequencing depth, method and number of samples were not sufficient, and therefore, further research is needed to validate this result. Another striking difference was the small number of chemosenory genes in PRM compared with other homologous species (Fig. 16), which possibly indicates differences in their ability to discriminate volatile chemical compounds and communicate with conspecifics. This difference may have evolved as a result of very different nesting behaviours, feeding habits and host-specificity of the poultry red mite. For example, most of endoparasitic mite species do not actively seek out their hosts. Transmission occurs through close or direct physical contact. The loss of active host-seeking behavior might thus have resulted in a reduction both in number of genes involved in chemosensation and in sensillum numbers. Both commonly observed in species that transition from ectoparasites to endoparasites.

**Fig. 16.**
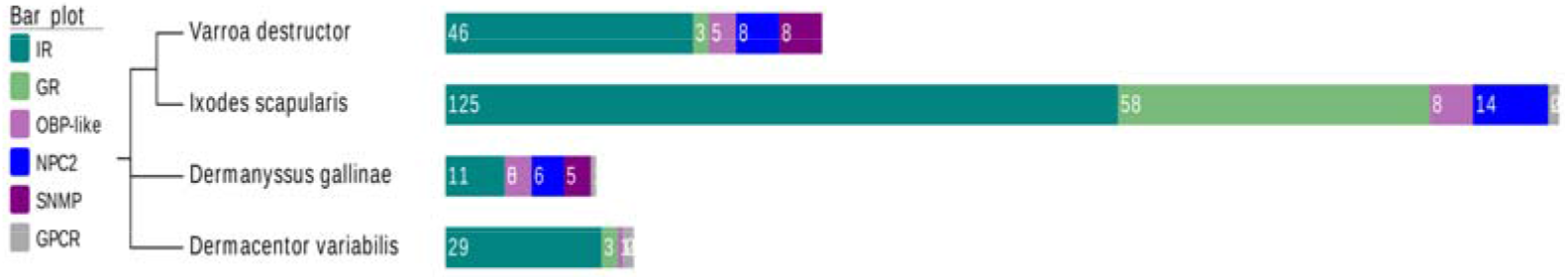
The number of chemosensory genes in different mite and tick species are presented in bar plot. A phylogenetic tree showing evolutionary relationships between various species are illustrated on the left. The number of chemosensory genes are obtained from *V. destructor, I. scapularis, D. variabilis*.

OBPs have recently been reported in some chelicerata, demonstrating that these proteins are structurally reminiscent of insect odorant binding proteins, and therefore named OBP-like. These proteins were first described in the tick *Amblyomma americanum* and the honey-bee mite *V. destructor*, suggesting a likely involvement in acari chemodetection (Renthal et al., 2017; Eliash et al., 2017). These secreted proteins have six cysteine amino acid residues as is also observed in the classical OBP family. The arrangement pattern of six conserved cysteines in the *D. gallinae* OBP family is comparable to the patterns in other OBP-like proteins of chelicerata (Fig. 7a). A major finding of our study is the identification of transcripts encoding OBP-like soluble proteins, some of which were highly and differentially expressed in the forelegs. This was also evident in *I. scapularis* ticks, where some of the OBP-like transcripts were highly expressed in the forelegs (Josek et al., 2018). Phylogenetic analysis revealed that all *Dermanyssus* OBPs were clustered within arachnida only, indicating that this branch evolved after the splitting of insects and arachnids. The continued annotation of OBP genes from chelicerate genomes will likely clarify this issue. In addition, soluble proteins of a different class (NPC2), which was also detected in the *varroa* proteomic project, have been suggested to function as semiochemical transporters in chelicerates (Pelosi et al., 2014; Renthal et al. 2017). This study found that NPC2 was expressed not only in chemosensory sensilla, but also in other parts of the body, suggesting roles in both chemical detection as well as in other functions. The results of this study were consistent with other studies in arthropods, which showed wide expression of NPC2 proteins across the body (Ishida et al., 2014; Pelosi et al., 2017; Zhu et al., 2018).

In the transcriptome analysis we also identified sensory neuron membrane proteins (SNMPs), which have key functions in pheromone detection in insects. The role of SNMPs in chemoreception in acari is still unknown. Similarly, there is little information available on the identified GPCRs, and the molecular characterization of most GPCRs remains unknown in acari. Identification of G proteins are also relevant for the understanding of acari biology in general, as these proteins could be targeted by novel compounds. Most information on acari GPCRs were obtained from tick transcriptome datasets and recombinant receptor systems (Gross et al., 2015; Xiong et al, 2019). Previous experiments in ticks indicated that prediction of GPCRs using transcriptome analysis might be a challenging task. As such, the transcriptome study of the bovine tick *Boophilus microplus* did not allow to detect the kinin receptor, a neuropeptide GPCR (Guerrero et al., 2016), which should be present (Holmes et al., 2000), underscoring the often low expression of GPCRs as a challenge for detection. Likewise, only two G proteins were identified in *I. scapularis* ticks and all were expressed at low to non-existent levels in legs (Josek et al., 2018). All of these findings acknowledge limitations in the prediction of tick GPCRs. In agreement with this, only one GPCR transcripts was detected in the present study. In addition, the presence of putative rhodopsin-like GPCR transcripts, having a foreleg bias expression in *Dermanyssus* mite, might code for olfaction. A similar finding was observed in the American dog tick *Dermacentor variabilis* with foreleg-biased expression of GPCRs, suggesting that tick olfaction via the Haller’s organ involves a GPCR-mediated pathway like that of vertebrates and nematodes (Carr et al., 2017). We thus suggest that mite olfactory receptors might represent a completely novel type of 7-transmembrane receptor (7TM) family proteins that have yet to be identified.

After identification of putative genes of interest, functional genomics tools such as RNA interference (RNAi) or CRISPR can be used to assess gene function. The RNAi approach has been employed widely to demonstrate knockdown of target genes, which subsequently validate the function of genes in order to identify novel candidate genes for drug or vaccine development (Palevich et al., 2018). This functional genomic approach can be applicable to other parasites, such as in mites where annotated functional genomic information is still limited (Gu and Knipple, 2013; Marr et al., 2014). A typical RNAi can be induced by a variety of delivery systems including noninvasive immersion, electroporation, micro-injection and feeding. The micro-injection alternative has widely been used for research in insects and ticks in spite of having high mortality due to trauma. However, the small size of mites (av. 900 × 510 μm) prevent micro-injection of dsRNA from being practical. Our present study demonstrated that a less invasive technique, using the non-invasive immersion method, can be used and is simple and less labor-intensive. Using the conserved odorant binding protein as a test case for RNA interference study, a significant reduction of gene transcript was achieved following 14 hours immersion in double stranded RNA encoding the OBP-1 gene. For efficient silencing, the dsRNA dosage and incubation at 4°C were chosen based on previous experiments (Marr et al., 2015), but this should be optimized for each future experiment. The development of gene silencing in PRM could be considered as a method of developing protocols for RNAi in other astigmatid mites of veterinary and medical importance.

## 6. Conclusions

The chemosensory genes reported here represent significant contribution to the pool of identified olfactory genes in non-insect arthropods in general. Mites consistently move their first pair of legs in the air in the manner of the antennae of insects, along with ultrastructural features of the sensory organs, and the results of transcriptomic analyses are all strongly in favour of the assumption that the forelegs serve an olfactory function. These results highlight the importance of an integrative approach in systematic studies, combining morphological and molecular characteristics, previously not being conducted in other mite and tick species. Further, the ability to induce RNAi through a simple non-invasive immersion-based dsRNA delivery in *D. gallinae* will allow researchers to study olfactory function in detail. Further work on other relevant genes needs to be done to verify these results.

## Acknowledgement

We sincerely thanks to Xie Yi (Chinese academy of tropical agricultural sciences, Hainan, China) for her help in SEM analysis. The authors thank Dr. Sunil Kumar Sahu (BGI-Shenzhen, China) and Jing Haiyu (BGI-Shenzhen, China) for RNA-seq analysis. We are grateful to Anindya Sen, Bijoy Sen, Kawser Ayon, Jing Chen, Lei Zhang, Tianlin Bi, Min Chen for their help in experiment and assistance with the figure production.

## Authors’ contributions

QH designed this study, obtained funding for this study, and supervised all research activities. BB drafted and edited the manuscript, and conducted experiments. YT, FL, JZ and CL performed bioinformatics analysis and figure production. ØØ, RI, BSH and QH commented on experimental design, methodology, and substantially revised the manuscript. All authors read and approved this final manuscript.

## Availability of data and materials

RNA-seq data discussed in this article are publicly available at the NCBI database under accession number PRJNA602095.

## Ethics approval and consent to participate

Not applicable.

## Consent for publication

Not applicable.

